# CD200R1 promotes IL-17 production by ILC3s, by enhancing STAT3 activation

**DOI:** 10.1101/2022.04.17.488595

**Authors:** Holly Linley, Alice Ogden, Shafqat Jaigirdar, Lucy Buckingham, Joshua Cox, Megan Priestley, Amy Saunders

## Abstract

Psoriasis is a common chronic inflammatory skin disease with no cure. It is driven by the IL-23/IL-17A axis and T_H_17 cells but, recently group 3 innate lymphoid cells (ILC3s) have also been implicated. However, the development, and factors regulating the activity of ILC3s remain incompletely understood.

Immune regulatory pathways are particularly important at barrier sites such as the skin, gut and lung, which are exposed to environmental substances and microbes. CD200R1 is an immune regulatory cell surface receptor which inhibits proinflammatory cytokine production in myeloid cells. CD200R1 is also highly expressed on ILCs, where its function remains largely unexplored. We previously observed reduced CD200R1 signalling in psoriasis skin, suggesting that dysregulation may promote disease. Here we show that contrary to this, psoriasis models are less severe in CD200R1-deficient mice due to reduced IL-17 production. Here we uncover a key cell-intrinsic role for CD200R1 in promoting IL-23-driven IL-17A production by ILC3s, by promoting STAT3 activation. CD200R1 is expressed on ILC precursors and is particularly high on neonatal ILC3s, suggesting CD200R1 may function during ILC development. Therefore, CD200R1 is required on ILC3s, potentially during their development, to promote IL-23-stimulated STAT3 activation triggering optimal IL-17 production.

## Introduction

Barrier sites such as the skin, are enriched for immune cells that can rapidly respond to stimulation by producing large quantities of proinflammatory cytokines. Innate lymphoid cells (ILCs) are one such cell type, with group 3 ILCs (ILC3s) responding to IL-23 and IL-1β by producing IL-17A and IL-22 (1). These cytokines cause changes in epidermal cells including hyperproliferation, dysregulated maturation, and epidermal production of cytokines, chemokines and antimicrobial peptides, which are critical for protection against extracellular bacterial and fungal infection (2, 3). However, the IL-23/IL-17 axis can also drive psoriasis, a common, incurable chronic inflammatory skin disease, as demonstrated by the efficacy of biologics targeting this pathway (4). ILC3s are also implicated in psoriasis as they are a potent source of IL-17 and are increased in lesions (5, 6). Therefore, it is crucial to understand how ILC3 activity is regulated to prevent inappropriate activation.

Immunosuppressive pathways are key to maintaining immune homeostasis and are particularly important in barrier tissues where there is constant exposure to environmental stimuli and commensal microbes. A key immunosuppressive pathway in myeloid cells is CD200R1 signalling, which has been shown to suppress proinflammatory cytokine production, reducing immune responses against self-antigens (7–10) allergens (11), infectious agents (12) and cancer (13). CD200R1 is a cell surface receptor which on engagement with its ligand, CD200, transmits a signalling cascade inhibiting MAP kinases and suppressing myeloid cell activation. CD200R1 is expressed on many immune cell types, and in addition to myeloid cells, is particularly highly expressed on ILC (14–17), however its function on these cells remains underexplored.

We have previously shown that the ligand, CD200 is reduced in non-lesional psoriatic skin (17) and we therefore hypothesized that in healthy skin, CD200R1 signalling prevents inappropriate inflammation, but in psoriasis this is reduced, contributing to psoriasis susceptibility. To examine this hypothesis, here we compare the severity of inflammation in psoriasis models in WT and CD200R1-deficient mice. This shows that surprisingly, global CD200R1 deficiency reduces the severity of psoriasis-like skin inflammation due to reduced IL-17A production. CD200R1 is highly expressed by ILCs and here we show that CD200R1-deficient ILC3s are less able to produce IL-17A. This is associated with a vastly different transcriptomic profile despite the surface phenotype being largely unchanged. We also demonstrate that the requirement for CD200R1 in ILC3 is cell intrinsic, but CD200R1 expression is highest on ILC3s during the neonatal period, suggesting that the role of CD200R1 may be during the development of these cells. To understand why CD200R1KO ILC3s are unable to produce IL-17A optimally, signalling downstream of IL-23 stimulation was investigated, demonstrating that CD200R1 is required for optimal STAT3 activation in response to IL-23. Therefore, here we describe a critical role for CD200R1 in promoting IL-17 production by ILC3 via enhancing IL-23-driven STAT3 activation.

## Results

### Global CD200R1 deficiency impairs IL-17 production in psoriasis models

As we (17) and others (18) have demonstrated reduced CD200 levels in psoriatic patient samples, and CD200R1 signalling has been shown to dampen immune responses (7–13), we hypothesized that the absence of CD200R1 would enhance inflammation in psoriasis models. Contrary to this, global CD200R1KO deficiency protected mice from skin thickening in a model induced by repeated intradermal injection with IL-23 (19) (Figure 1A and B). Total immune cells (CD45^+^) were increased in IL-23 treated WT mouse skin, but this increase was inhibited in CD200R1KO mice (Figure 1C). To determine how CD200R1 promotes inflammation, skin IL-17A levels were measured, as this is a critical cytokine driving inflammation. ELISA and flow cytometry demonstrated reduced IL-17A overall in skin, and reduced production by both ILCs and CD3^low^ γδ T cells in CD200R1KO mice. (Figure 1D-F) To investigate this further, a more complex psoriasis model was used in which Aldara cream, (containing the TLR7-agonist, imiquimod, and inflammasome activating isosteric acid) induces inflammation (20). After 4 days of treatment skin thickening was similar in WT and CD200R1KO mice, but redness and scaling were slightly reduced in the absence of CD200R1 (Figure 1G-I). The absence of CD200R1 reduced the accumulation of immune cells, in particular neutrophils, within skin (Fig 1J and K) and reduced the production of IL-17A by ILCs and γδ T cells within skin and LN (Figures 1L and M).

**Figure 1:**
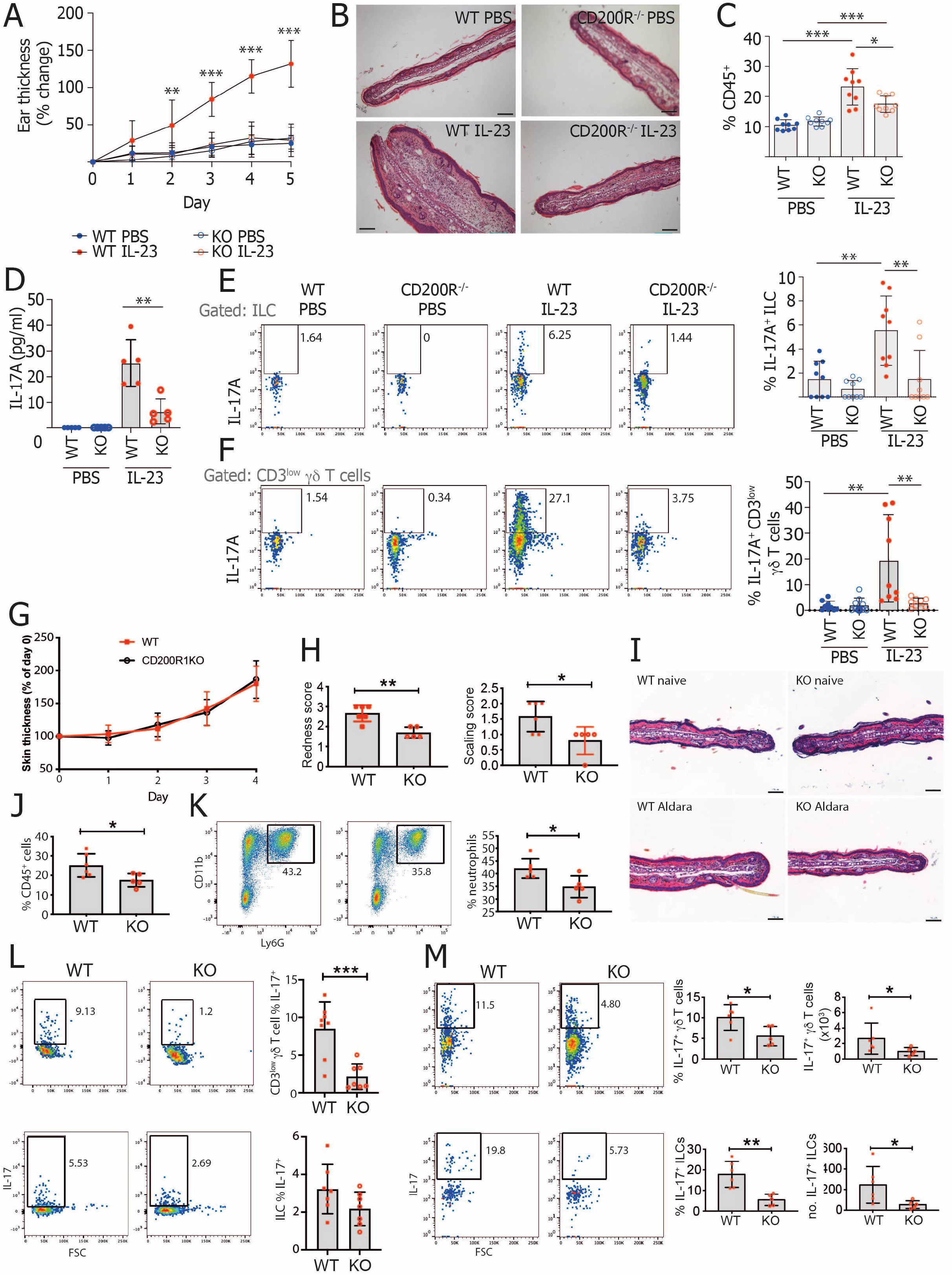
CD200R1 promotes IL-17A production in murine models of psoriasis. **A-F** WT and CD200R1KO (KO) Ears were intradermally injected with 1μg IL-23, or PBS daily for 5 consecutive days. **A.** Skin thickening measured using a digital micrometer. **B.** H&E stained skin sections. **C.** Proportion of live, single cells in skin that are CD45^+^. **D.** IL-17A level in media from overnight cultured skin cells. **E.** Proportion of skin ILCs (CD45^+^ Lin^-^ CD3^-^ TCRβ^-^ γδTCR^-^ Thy1^+^ CD127^+^) producing IL-17A. **F.** Proportion of skin CD3^low^ γδ T cells producing IL-17A. **G-M.** Inflammation was induced by topically treating WT and CD200R1KO (KO) ear skin with Aldara cream daily. **G.** Skin thickening measured using a digital micrometer. **H.** Skin redness and scaling scores. **I.** H&E stained skin sections. **J.** Proportion of live, single cells in skin that are CD45^+^. **K.** Proportion of skin CD45^+^ cells that are neutrophils (CD45^+^ CD11b^+^ Ly6G^+^). **L.** Proportion of skin CD3^low^ γδ T cells or ILCs producing IL-17A. **M.** Proportion of lymph node CD3^low^ γδ T cells or ILCs producing IL-17A. Histology scale bars show 100 μm. Data shown are pooled from 2 independent experiments except for B and I showing representative histology, D showing representative ELISA, H, J, K and M showing representative data from one experiment. A, C-F were analysed by two-way ANOVA followed by a Bonferroni post hoc test. H, J-M were analysed by Student’s t test.

IL-17 is critical to combatting fungal infections therefore, to examine if CD200R1KO mice also have altered fungal immune responses in skin, a *C. albicans* infection model was performed which showed CD200R1KO mice produce less IL-17 in response to fungal skin infection (Figure S1).

### CD200R1 is highly expressed by ILCs

Given the reduction in IL-17A production in psoriasis-like skin inflammation models, the expression of CD200R1 on skin cells was examined. As anticipated from the literature, CD200R1 was expressed on skin macrophage and dendritic cell subsets (gated as described previously (21)) but was absent from non-immune cells and dendritic epidermal T cells (DETC) (Figure 2A-D). CD200R1 was also expressed on a small proportion of dermal γδ T cells and on a larger proportion of skin ILCs (Figure 2D) as previously described (17). On splitting the ILCs into subsets based on their expression of lineage defining transcription factors, CD200R1 was shown to be expressed most highly on ILC2 but, was also expressed on ILC3s and GATA3^-^ RORγt^-^ ILCs in skin (Figure 2E). Given the low level of expression of CD200R1 on dermal γδ T cells, and the higher expression on ILCs, the effect of CD200R1 on ILCs was examined.

**Figure 2:**
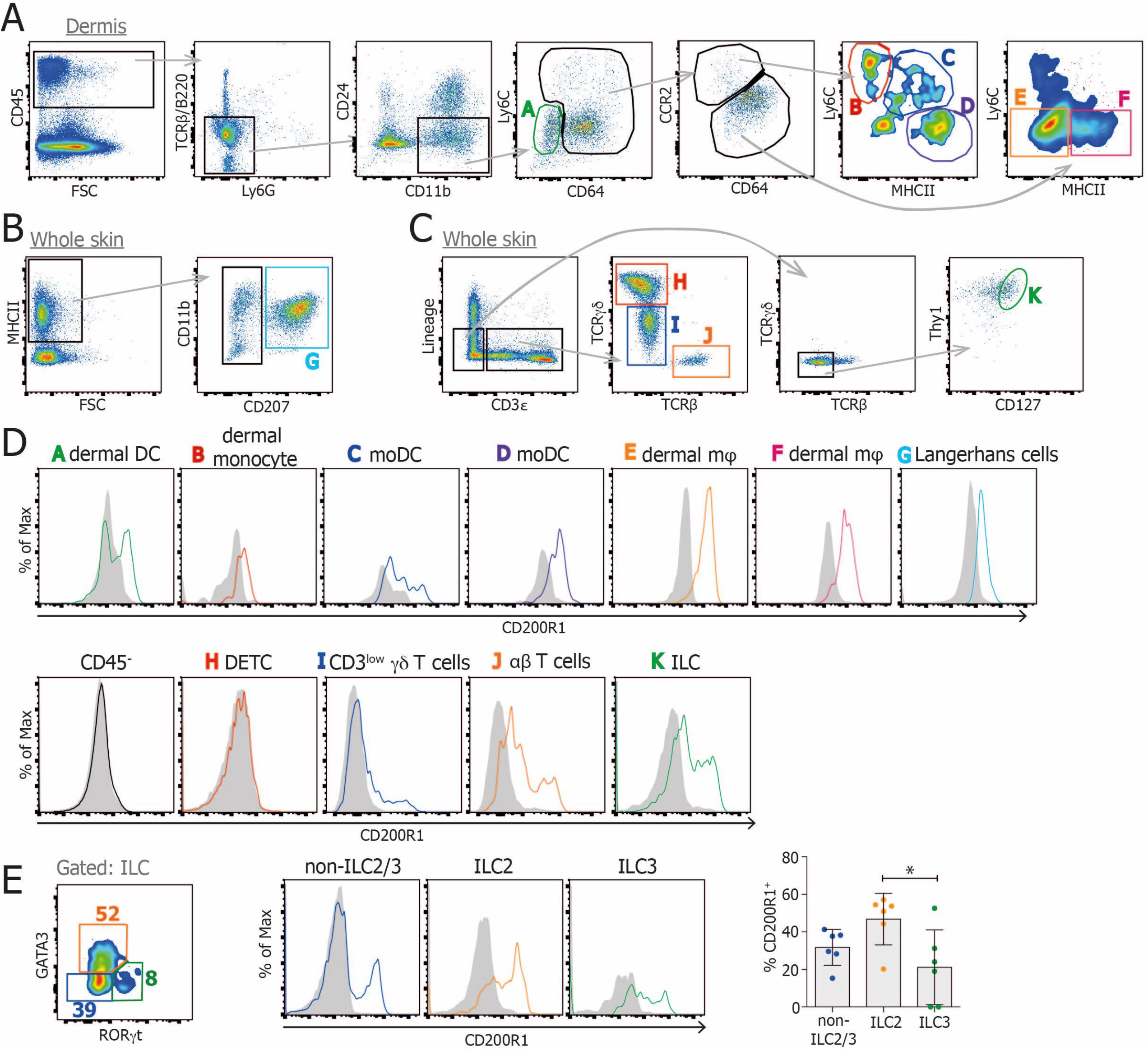
CD200R1 is expressed by skin ILCs. Dorsal skin cells were isolated and analysed by flow cytometry. **A.** Dermal live CD45^+^ single cells were gated into various monocyte, dendritic cell and macrophage populations. **B.** Whole skin live CD45^+^ single cells were gated for Langerhans cells. **C.** Whole skin live CD45^+^ single cells were gated for T cell subsets and ILCs. **D.** CD200R1 expression on each cell subset. **E.** Skin ILCs (CD45^+^ Lin^-^ CD3^-^ TCRβ^-^ γδTCR^-^ Thy1^+^ CD127^+^) were gated for ILC2 (GATA3^+^), ILC3 (RORγt^+^) and non-ILC2/3 (GATA3^-^ RORγt^-^), and CD200R1 expression was analysed within each subset. Filled histograms show CD200R1KO data, lines show WT data. All data are representative of 2 replicate experiments except E showing data from 2 experiments which was analysed by One-way ANOVA.

### CD200R1 promotes IL-17A production by ILCs

To determine if CD200R1 regulates IL-17 production by ILCs in a more direct manner, outside of the context of skin inflammation, dorsal skin cells were isolated and stimulated with IL-23, or PMA and Ionomycin, demonstrating that CD200R1-deficient ILCs are less able to produce IL-17A than WT, regardless of the stimulatory agents used (Figure 3A and B). IL-22 production was also examined and may be reduced in CD200R1KO ILCs, but this was not found to be statistically significantly different, potentially due to the relatively low levels of production (data not shown). Similarly, gut ILCs from CD200R1-deficient mice were less able to produce IL-17A after stimulation with PMA, Ionomycin and a cocktail of cytokines, regardless of whether they are cultured unseparated, or cultured after flow cytometric sorting to obtain a purified ILC population (Figure 3C). This reduction in IL-17A production by CD200R1 deficient ILCs was found to be independent of secreted inhibitory factors in the CD200R1-deficient cell cultures, as co-culture of WT and CD200R1-deficient cells maintained the impaired IL-17A production by CD200R1KO ILCs (Figure 3D). There was also no increase in known ILC3 inhibitory cytokines, IL-10, IFNγ and IL-25 in the CD200R1-deficient cell cultures relative to WT cultures (Figure S2), suggesting that CD200R1 directly affects IL-17 production rather than modulating the activity of an intermediary cell type in this *in vitro* model.

**Figure 3:**
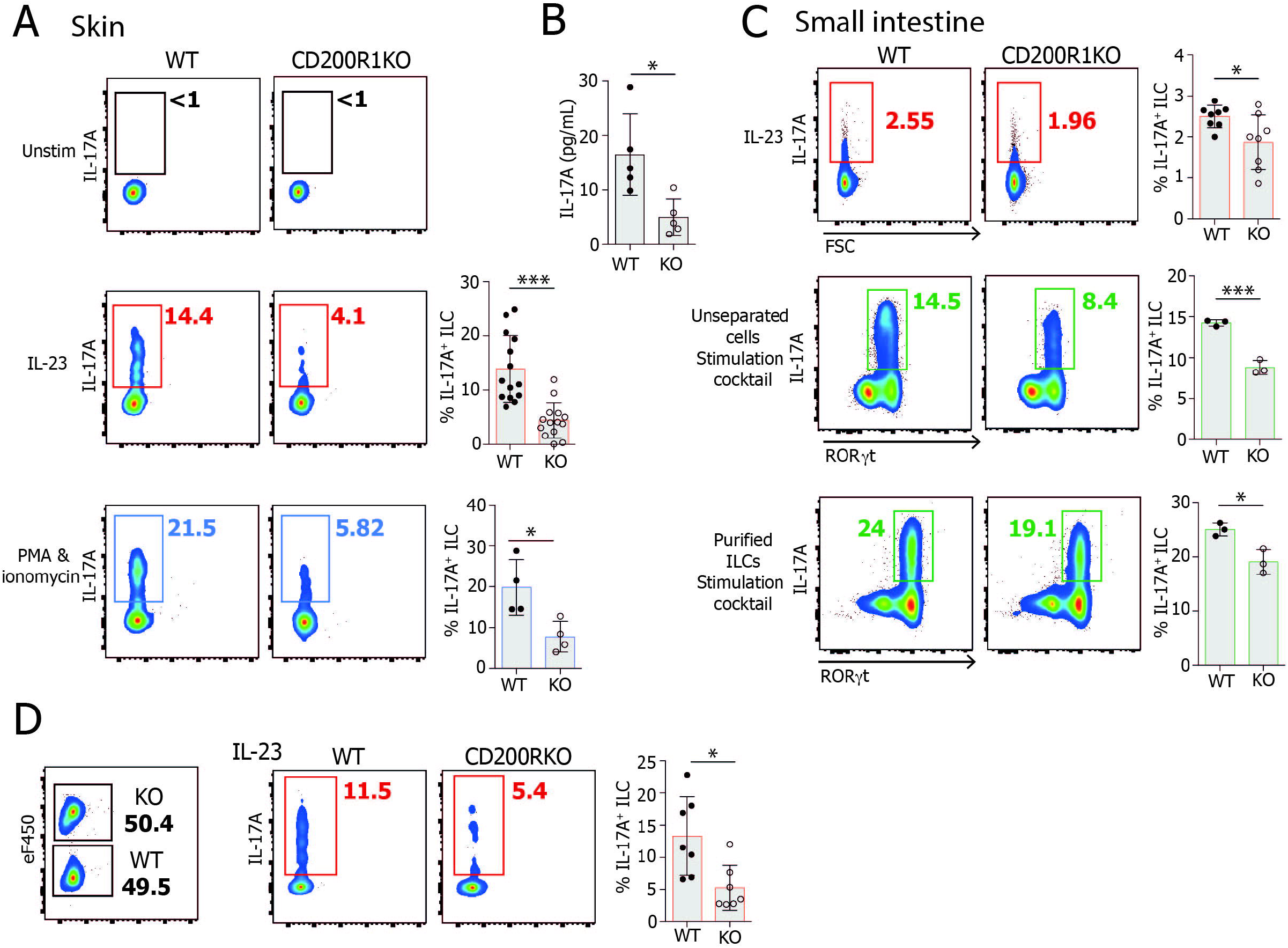
CD200R1 is required for efficient IL-17A production by ILCs. Dorsal skin or intestinal lamina propria cells from WT or CD200R1KO (KO) mice were stimulated in culture and IL-17A production by ILC was measured by flow cytometry. **A.** IL-17 production by dorsal skin ILCs (CD45^+^ Lin^-^ CD3^-^ TCRβ^-^ γδTCR^-^ Thy1^+^ CD127^+^) stimulated with IL-23, or with PMA and ionomycin for 4 hrs. **B.** IL-17A levels in cell supernatant from IL-23-stimulated skin cells. **C.** IL-17 production by intestinal lamina propria cells, or flow cytometric sorted intestinal lamina propria ILCs (CD45^+^ Lin^-^ CD3^-^ TCRβ^-^ γδTCR^-^Thy1^+^ CD127^+^) stimulated with IL-23 alone or PMA, Ionomycin, IL-23, IL-6, IL-2 and IL-1β for 5 hrs. **D** CD200R1KO dorsal skin cells were labelled with a fluorescent dye before being co-cultured in a 1:1 ratio with unlabelled WT cells. IL-17A production was analysed in co-cultured cells stimulated with IL-23 overnight. A (PMA/Ionomycin stim), B, C (stimulation cocktail) data shown are from one representative experiment. C (IL-23 stim), D data shown are from two independent experiments. A (IL-23 stim) data shown are from 3 independent experiments. Data were analysed by Student’s t test.

### CD200R1 does not affect ILC3 subsets

We hypothesized that the reduced IL-17 production seen in CD200R1KO ILCs may be due to either a reduction in ILC3s numbers, or the lack of an ILC3 subset. To test this hypothesis, firstly the proportion and numbers of ILC2, ILC3 and ILC lacking GATA3 and RORγt (which will contain ILC1 and ILCs which have lost expression of lineage defining transcription factors (22)) were examined in skin and small intestinal lamina propria. This showed there was not a reduction in ILC3 in CD200R1KO which could account for the reduced IL-17 production (Figure 4A-B). ILC3 are heterogeneous, particularly in the small intestine where 4 main subsets of ILC3s are seen (23). NCR^+^ (NKp46^+^) ILC3s are specialised in interacting with myeloid and stromal cells, CD4^+^ and CD4^-^ Lti-like (CCR6^+^) ILC3s have roles in modulating innate and adaptive immune responses, and NCR^-^ ILC3s are precursors of the NCR^+^ ILC3 subset. All of these subsets were observed in CD200R1KO small intestine at similar frequencies to WT mice (Figure 4C-D), showing the reduced IL-17 production in CD200R1KO ILCs is not due to a missing major ILC3 subset. Less is known about the subsets of ILC3 present in skin but, applying similar gating as that used for the small intestine allowed the presence of CD4^-^ Lti-like ILC3s and NCR^-^ ILC3s to be observed. Similar to the small intestine, CD200R1 deficiency did not affect these skin ILC3 populations (Figure 4E-F).

**Figure 4:**
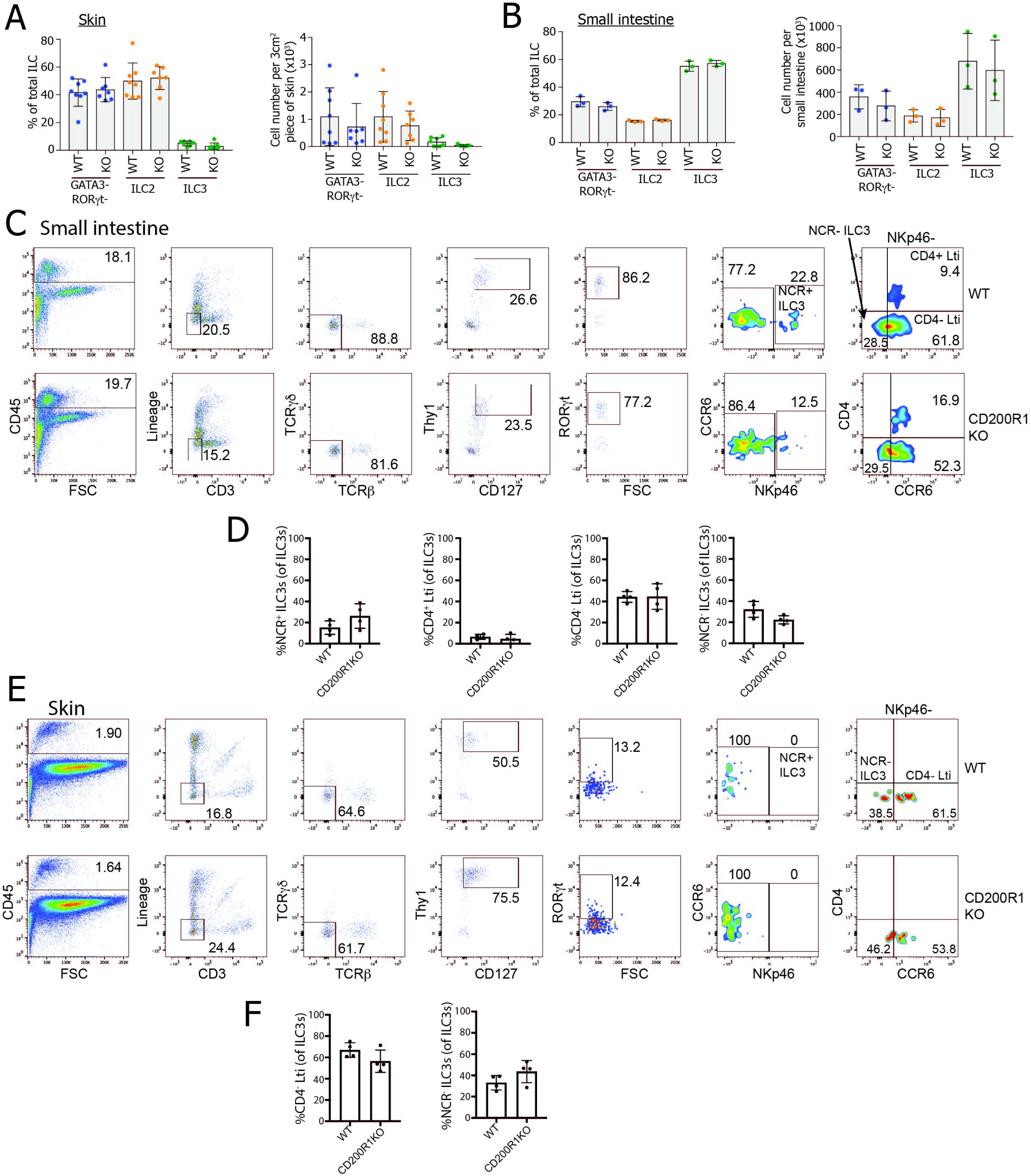
CD200R1KO mice do not lack any major subsets of ILC3s. **A.** The proportion and number of WT and CD200R1KO (KO) skin ILCs (CD45^+^ Lin^-^ CD3^-^ TCRβ^-^ γδTCR^-^ Thy1^+^ CD127^+^) within each subset (ILC2, GATA3^+^; ILC3 RORγt^+^; non ILC2/3, GATA3^-^ RORγt^-^). **B.** The proportion and number of small intestinal lamina propria ILCs within each subset. **C.** The proportion of ILC3s in each subset in WT and CD200R1KO small intestinal lamina propria. **D.** The proportion of ILC3s in each subset in WT and CD200R1KO skin. A data shown from 2 independent experiments. B-F data are representative of 2 independent experiments.

### CD200R1-deficient ILC3s are transcriptomically very different to WT

To determine why CD200R1-deficient ILC3s are less able to produce IL-17A, the cell surface phenotype was analysed by flow cytometry. Skin ILC3s showed undetectable CD4 and ckit and only low levels of CD25, IL-23R and CD1d expression. They expressed CCR6, Sca-1 and MHC class II, but CD200R1 deficiency does not appear to affect the expression of any of these markers (Figure 5A). In the gut, expression of all of the markers was observed, but again, CD200R1 deficiency did not affect the levels of these markers (Figure 5B), demonstrating that CD200R1-deficient ILC3 have a similar cell surface phenotype to WT. Gut ILC3 are highly heterogeneous, so there may be differences in the expression of some of these markers on a specific ILC3 subsets. However, as the main ILC3 subsets are in equal proportions in CD200R1KO and WT mice (Figure 4B-D), this seems unlikely. To determine how CD200R1 affects ILC3 function, ILC3s from WT (RORc eGFP) and CD200R1-deficient (CD200R1KO RORc eGFP) mice were sorted from small intestine (total ILC3s isolated, gating and purity shown in Figure S3) and RNASeq was performed. Small intestinal cells were used to provide a large enough cell population to be isolated. CD200R1KO ILC3s were found to have 3285 differentially regulated genes, with 2252 of those upregulated, and 1033 down-regulated relative to WT (Figure 5C). Despite there being more upregulated than down regulated genes in the CD200R1-deficient ILC3s, there are less highly upregulated genes (with a log2 fold change of >10) than there are highly down regulated genes (with a log2 fold change <-10) (Figure 5D). Pathway analysis revealed many changes in CD200R1-deficient ILC3s, but the most highly enriched pathway was the nuclear factor-erythroid 2–related factor 2 (NRF2)-mediated oxidative stress response (Figure 5E-F), which is known to protect against oxidative stress and reduce ROS production but also inhibits Th17 differentiation, STAT3 activation and innate immune cell activation (24, 25). Indeed, other pathways enriched in CD200R1KO ILC3s included STAT3 signalling as well as others of potential interest including TGFβ signalling, and embryonic stem cell pluripotency (Figure 5E-F). A small number of pathways were down-regulated in CD200R1KO ILC3, the most significantly altered being interferon signalling (Figure 5G-H). This may reflect an inability to signal in response to interferons, or a reduction in interferons in CD200R1KO mice. Overall these data suggest that CD200R1-deficient ILC3s are transcriptomically very different from WT and may have changes in cytokine signalling pathways.

**Figure 5:**
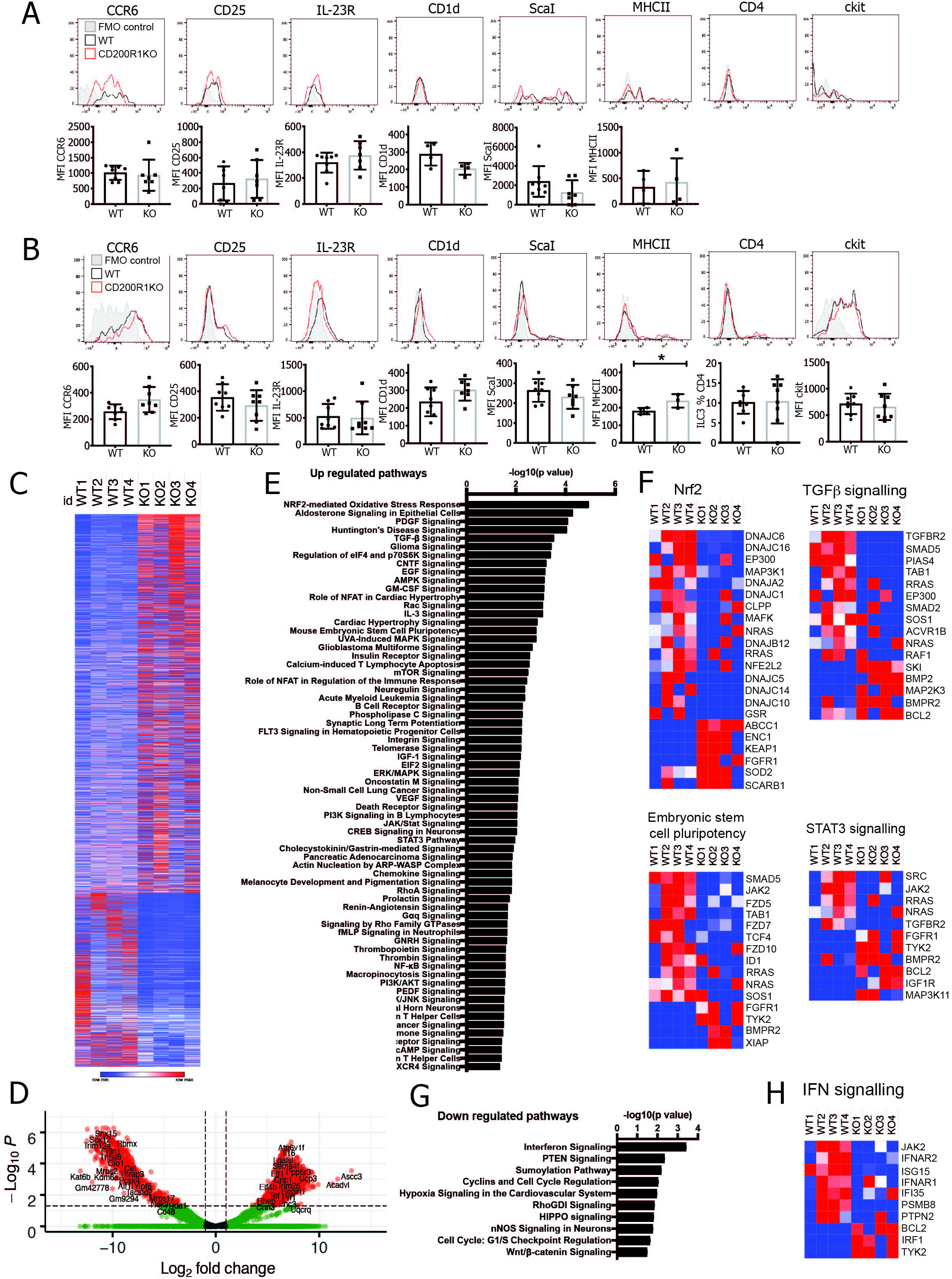
CD200R1KO ILC3s are deficient at activating STAT3. WT and CD200R1KO (KO) ILC3s (CD45^+^ Lin^-^ CD3^-^ TCRβ^-^ γδTCR^-^ Thy1^+^ CD127^+^ RORγt^+^) cell surface marker expression in **A.** dorsal skin, **B.** small intestinal lamina propria. **C.** RNAseq analysis of flow cytometrically sorted ILC3s from WT (RORγt eGFP) and CD200R1KO (CD200R1KO RORγeGFP) lamina propria. Log2fold change values are shown for all differentially regulated genes. **D.** Volcano plot of the RNAseq data. **E.** Pathways predicted to be upregulated in KO ILC3 by pathway analysis of the RNAseq data. **F.** Heat maps showing z scores for genes in pathways predicted to be enriched in KO ILC3. **G.** Pathways predicted to be reduced in KO ILC3 by pathway analysis of the RNAseq data. **H.** Heat maps showing z scores for genes in pathways predicted to be downregulated in KO ILC3. A, B. data are from 2 independent experiments. Data were analysed by Student’s t test.

### CD200R1 plays a cell intrinsic role in promoting IL-17A production by ILC3s

Previously we showed that blocking CD200R1 in adult mice did not affect IL-17 production by ILCs (17), however ILCs seed tissues during development, so a global deficiency in CD200R1 may have earlier developmental effects. In support of this, embryonic stem cell pluripotency genes were found to be increased in CD200R1KO ILC3s (Figure 5E-F). To determine if CD200R1 is differentially expressed with age, CD200R1 levels in ILC3s from mice of different ages were examined. CD200R1 is particularly highly expressed by ILC3s from neonatal mice but this is reduced between the neonatal period and weaning (3 weeks old) (Figure 6A). To examine potential defects in ILC progenitors, WT and CD200R1KO bone marrow cells were examined (Figure 6B-D). CD200R1 is expressed on ILC progenitors, particularly immature ILC2, but also at lower levels on common helper-like innate lymphoid progenitor (CHILP) and ILC precursor populations (Figure 6C). Although immature ILC2 populations are unchanged in CD200R1KO mice, ILC precursors are increased and CHILP are decreased, suggesting that CD200R1 effects the development of ILCs (Figure 6D).

**Figure 6:**
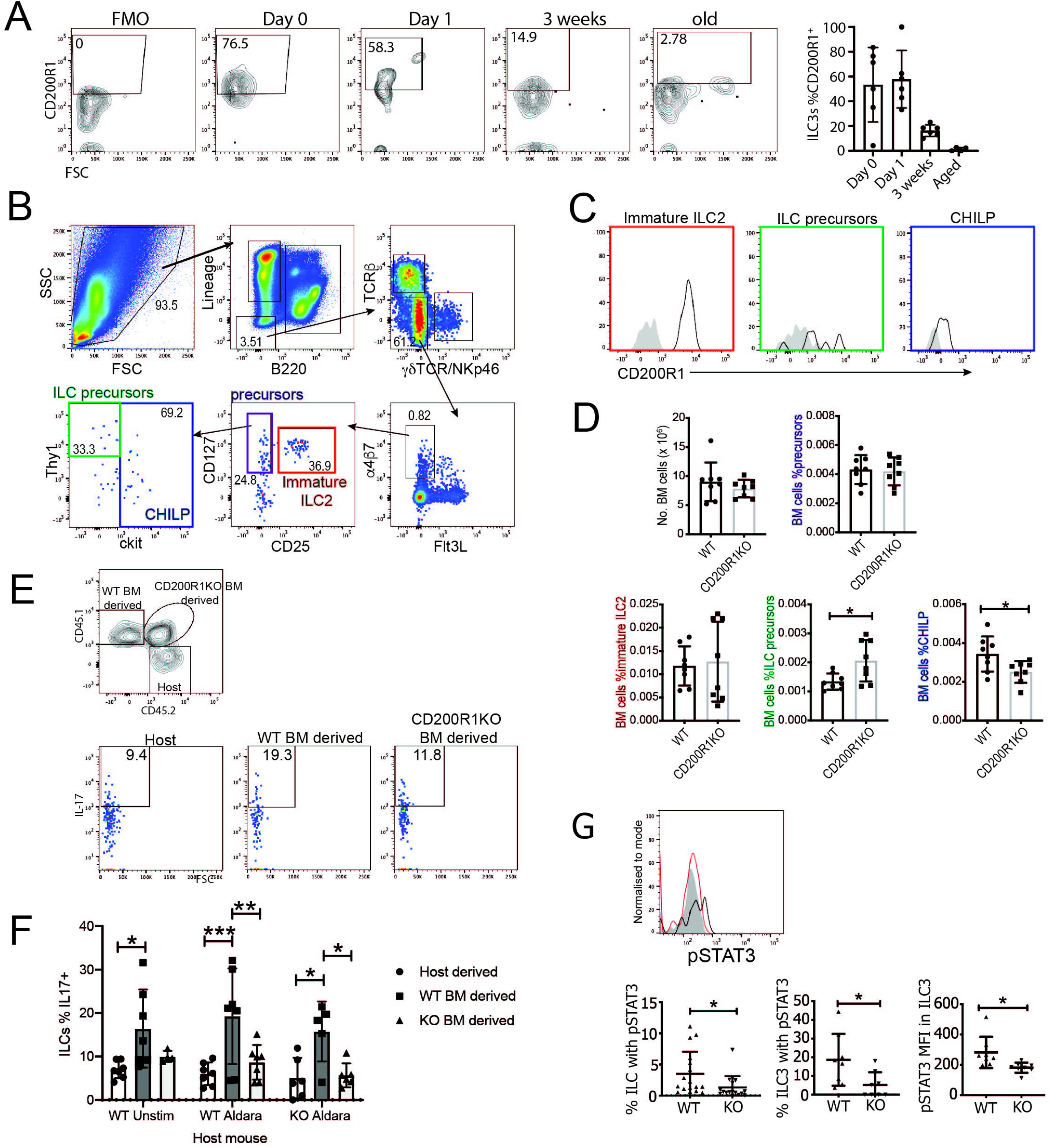
CD200R1 is required in a cell intrinsic manner to licence ILC3 for IL-17A production. **A.** CD200R1 expression on dorsal skin ILC3s (CD45^+^ Lin^-^ CD3^-^ TCRβ^-^ γδTCR^-^ Thy1^+^ CD127^+^ RORγt^+^) from 0 or 1 day old, 3 week-old and partially aged (1 year old) mice. **B.** Gating strategy for ILC progenitors in adult bone marrow. **C.** CD200R1 expression on ILC progenitor subsets in adult bone marrow. **D.** ILC progenitor cell populations in WT and CD200R1KO bone marrow. **E.** Representative plots showing IL-17 production by ILC in mixed bone marrow (WT and CD200R1KO) chimeric mice. **F.** All data showing IL-17 production by ILC in mixed bone marrow (WT and CD200R1KO) chimeric mice. **G.** Proportion of pSTAT3 positive ILCs and ILC3s, and median fluorescence intensity of pSTAT3 in lymph node ILC3 stimulated with IL-23 for 20 min. Representative histogram showing pSTAT3 staining on WT ILC3s (black line), CD200R1KO ILC3s (red line) or pSTAT3 FMO control (grey filled), and bar charts showing all data. A, D-G (ILC3 data) data are from 2 independent experiments. G (total ILC data) are from 3 independent experiments. D, G data were analysed by Student’s t test. F data were analysed by two-way ANOVA followed by a Bonferroni post hoc test.

To determine if CD200R1 affects ILC3 activity in a cell intrinsic or extrinsic manner, chimeric mice were generated with a mixture of WT and CD200R1KO bone marrow, and skin inflammation was induced. This revealed that CD200R1KO ILC are less able to produce IL-17A than WT ILCs in skin (Figure 6E-F), showing that CD200R1 promotes IL-17 production by skin ILC3s in a cell intrinsic manner.

### CD200R1 promotes ILC3 IL-17 production by enhancing STAT3 activation in response to IL-23

As the transcriptomic data showed that CD200R1KO ILC3 were enriched for cytokine signalling transcripts (Figure 5E-H), and CD200R1KO ILC3s are less able to produce IL-17A in response to IL-23 but have normal IL-23R levels (Figure 5A-B), we considered cytokine signalling downstream of IL-23R. STAT3 is a crucial mediator of IL-23 signalling, and the STAT3 signalling pathway was found to be enriched in CD200R1KO ILC3 (Figure 5E-F). This enrichment in signalling components may suggest there is an enhanced response to IL-23, or alternatively it may reflect the cell compensating for a deficiency in signalling. On engagement of IL-23R, Jak2 and Tyk2 become activated and phosphorylate STAT3 allowing its dimerization and translocation to the nucleus where it can modulate transcription of target genes (26). The RNAseq data suggest that STAT3 signalling may be affected by CD200R1 (Figure 5E), with increased Tyk2, but decreased Jak2 transcripts in CD200R1KO ILC3 (Figure 5F), suggesting that STAT3 signalling may be altered in CD200R1-deficient ILC3s. Therefore, the ability of CD200R1KO ILC3 to signal in response to IL-23 was analysed by measuring phosphorylated STAT3 (pSTAT3) levels. This demonstrated the proportion of pSTAT3 positive cells and the level of pSTAT3 is reduced in CD200R1-deficient ILCs and ILC3s (Figure 6G), showing that signalling in response to IL-23 is impaired in the absence of CD200R1.

## Discussion

Here we show that CD200R1 promotes IL-17A production by ILCs via the promotion of STAT3 phosphorylation. This is an intrinsic requirement for CD200R1 which likely manifests during ILC3 development or tissue seeding. This is surprising as the CD200:CD200R1 pathway has been shown to inhibit proinflammatory cytokine production by myeloid cells (27, 28), and suppress a variety of immune responses (7–13). However, most studies examining the role of CD200:CD200R1 signalling used exogenous ligand or blocked CD200R1 using antibodies. Therefore, the role of this pathway during the development of immune cells in the neonatal period has not previously been fully addressed.

We recently examined psoriasis-like skin inflammation in mice treated with a CD200R1 blocking antibody and found that IL-17A production was not changed, but neutrophil accumulation was increased, demonstrating that CD200R1 limits neutrophil recruitment (17). Here, we did not observe effects on neutrophil recruitment in the IL-23 injection model (data not shown), and only saw a slight decrease in neutrophils in the Aldara cream-induced psoriasis model. However, neutrophil recruitment effects may be masked in CD200R1-deficient mice due to compensation between the reduction in IL-17A and an increase in CD200R1KO neutrophil recruitment. Indeed, the skin thickening in response to psoriasis-like inflammation was not changed in CD200R1-deficient mice despite the reduction in IL-17A, which suggests that there may be an increase in an alternative inflammatory response in these mice. The fact that we did not observe a decrease in IL-17A production in psoriasis-like skin inflammation when CD200R1 was blocked in adult skin (17), supports our findings here that the role for CD200R1 promoting IL-17A production, is during an earlier development stage, perhaps in neonates when ILCs develop and seed peripheral tissues.

Other work has shown that mice genetically lacking the ligand, CD200, have reduced suppression, including in influenza and mouse coronavirus infections (12, 29, 30), in an experimental meningococcal septicaemia model (31) and in experimental autoimmune encephalitis and collagen induced arthritis models (32). This suggests that the presence of endogenous CD200 during development and adulthood does indeed inhibit diverse immune responses, which is opposite to the effects that we describe here for its receptor, CD200R1. However, recent papers suggest that the CD200:CD200R1 signalling pathway may not act as a straightforward ligand-receptor pair. The effect of exogenous CD200 is not mirrored by the effect of agonistic antibodies against CD200R1 (33), suggesting that either the agonistic antibodies do not activate CD200R1 in the same way as CD200, or that CD200 has effects independent from CD200R1. Although there exists CD200R-like proteins which appear to have activating functions and could account for differential effects of CD200, these do not appear to be capable of CD200 binding (34, 35), so probably do not account for these effects. Similarly, viruses are known to express CD200 homologues which are proposed to allow viral evasion of the immune response (36), however it has been shown that some of these effects are independent of CD200R1 (37). This again suggests that CD200 has effects independent of CD200R1. Conversely, in the gut other ligands for CD200R1 have been identified (38), and it is not yet clear if these ligands play similar, or distinct roles to CD200. In support of our finding here that CD200R1 can promote inflammation, a recent report, suggests that CD200R1 switches from being suppressive to being stimulatory in the presence of IFNα (39). Therefore, it is clear that we do not yet fully understand the different roles played by CD200 and CD200R1, and how these effects are mediated. Gaining a better understanding of this pathway is crucial, particularly as therapeutics are being developed to target this pathway for the treatment of cancers (40).

It is apparent that the factors in early life which determine how sensitive an individual is to immune challenges, and the types of immune response that are generated are not well understood. Here we show that CD200R1 is more highly expressed by ILC3s in early life, which suggests a role for CD200R1 in the development or differentiation of these cells. In the absence of CD200R1, STAT3 activation in response to IL-23 stimulation is blunted which leads to reduced IL-17 production.

One interesting finding here, is that CD200R1 also promotes ILC3 IL-17A production in response to PMA/Ionomycin stimulation, which bypasses the requirement for STAT3 activation. Whilst we do not yet understand the mechanism by which CD200R1 acts, given the large transcriptomic differences, it seems likely that CD200R1-deficiency dysregulates signalling pathway components more widely than just the STAT3 signalling pathway.

In conclusion, here was show a crucial novel role for CD200R1 in promoting IL-17A production by ILC3s, perhaps during the development/differentiation of these cells.

## Materials and Methods

### Mice

All animal experiments were locally ethically approved and performed in accordance with the UK Home Office Animals (Scientific Procedures) Act 1986. C57BL/6 mice were obtained from Charles River Laboratories. CD200R1KO mice (41), (provided by Prof. Tracy Hussell), RORc eGFP(42), (provided by Prof. Gerard Eberl), CD45.1 (43) (B.6SJL-Ptprca Pep3b/BoyJ–Ly5.1 mice, provided by Prof. Andrew MacDonald), CD200R1KO CD45.1 and CD200R1KO RORc eGFP mice (generated by crossing CD200R1KO with CD45.1 or RORc eGFP mice respectively) were bred and maintained in SPF conditions in house. Mice were 7 to 14 weeks old at the start of procedures, unless otherwise stated.

### Skin inflammation models

Mice were anaesthetised and ear thickness was measured daily using a digital micrometer (Mitutoyo). Redness and scaling were scored daily. For the IL-23 intradermal injection model, ears were injected intradermally with 1 μg IL-23 (eBioscience) or PBS, daily for five consecutive days (19). For the Aldara cream model (20), ears were treated topically with 20 mg Aldara cream (Meda Pharmaceuticals), (containing imiquimod and isosteric acid) daily, for 4 days.

### H&E staining

Ear tissue was fixed in 10% neutral buffered formalin then embedded in paraffin and cut to 5 μm. Haematoxylin and eosin staining was carried out using an automated Shandon Varistain V24-4. Images were acquired using a 3D-Histech Pannoramic-250 microscope slide-scanner. Snapshots were taken with Case Viewer software (3D-Histech).

### Cell isolation

#### Skin and LN

Ears were split in half and floated on 0.8% w/v Trypsin (Sigma) at 37 °C for 30 min, then epidermis was removed and both epidermis and dermis were chopped and digested in 0.1 mg/ml (0.5 Wunch units/ml) Liberase TM (Roche) at 37 °C for 1 hr. Subcutaneous fat was removed from dorsal skin before floating on Trypsin as above. Tissue was chopped and digested with 1 mg/ml Dispase II (Roche) at 37 °C for 1 hr. Digested skin tissue suspensions, or auricular lymph nodes were passed through 70 μm cell strainers and cells were counted.

#### Small intestinal lamina propria cells

Cells were isolated as detailed previously (44). Briefly, small intestines were excised and Peyer’s patches were removed. Intestines were sliced open, washed with PBS and cut into segments and shaken in RPMI with 3% FCS, 20mM HEPES and 1% penicillin streptomycin. The tissue was transferred to pre-warmed RPMI supplemented with 3% FCS, 20mM HEPES, 1% penicillin streptomycin, 2mM EDTA (Fischer Scientific) and 14.5 mg/mL dithiothreitol (DTT; Sigma) and incubated at 37°C for 20 minutes with agitation. The tissue was shaken 3 times in pre-warmed RPMI supplemented with 20 mM HEPES, 1% penicillin streptomycin and 2mM EDTA then was washed with PBS. Tissues were minced then digested with 0.1 mg/mL Liberase TL (Roche) and 0.5 mg/mL DNaseI in RPMI for 30 minutes at 37°C. Cell suspensions were passed through 70 μm, and 40 μm cell strainers (Fischer Scientific) then washed in complete RPMI (RPMI supplemented with 10% heat inactivated FBS, 1% penicillin streptomycin solution, 2 mM L-glutamine, 1 mM sodium pyruvate, 20 mM HEPES, 1X non-essential amino acid solution, 25 nM 2-mercaptoethanol (Sigma)).

#### Bone marrow cells

Fibulas and tibias were removed, and bone marrow cells were isolated using a needle and syringe and PBS. Red blood cells were lysed using ACK lysis buffer (Lonza) and cells were washed and counted.

#### Flow cytometric analysis

Cells were incubated with 0.5 μg/ml anti-CD16/32 (2.4G2, BD Bioscience) and Near IR Dead cell stain (Invitrogen) prior to staining with fluorescently labelled antibodies. Cells were fixed with Foxp3/Transcription Factor Buffer Staining Set (eBioscience) for between 30 min and 16 hr at 4°C.

For cytokine analysis, 10 μM Brefeldin A was added to cell cultures for 4 hr prior to staining for cell surface markers as described above. After overnight fixation, cells were permeabilized with Foxp3/Transcription Factor Buffer Staining Set (eBioscience) and were stained with intracellular markers or cytokines. Cells were analysed on a BD Fortessa or LSRII flow cytometer. Data were analysed using FlowJo (TreeStar). Antibodies are detailed in Table S1.

For pSTAT3 staining, LN cells were rested in culture in RPMI with 2% FBS for 1 hr, before stimulation with 40 ng/ml IL-23 (Biolegend) for 20 min. Cells were stained for surface markers, then fixed with Phosflow Fix buffer I (BD Biosciences) at 37 °C for 10 min, before fixation overnight at 4 °C in Foxp3/Transcription Factor Buffer Staining Set (eBioscience). Cells were incubated at 4 °C in Phosflow Perm Buffer III (BD Biosciences) for 30 min, then were stained with PE conjugated pSTAT3 (pY705) (BD Bioscience clone 4/P-STAT3).

ILCs were gated as live, single cell, CD45^+^, Lin^-^ (Ter119, F4/80, CD11b, CD11c, FceRIa, Gr1, CD19), CD3^-^, TCRb^-^, TCRgd^-^, CD90.2^+^ CD127^+^.

#### *In vitro* ILC activation assay

Mouse dorsal skin cells were stimulated in culture with 40 ng/mL IL-23 (Biolegend) alone, or in combination with 40 ng/mL IL-1β (eBioscience) overnight. Supernatants were retained for cytokine analysis and cells were cultured with 10 μM Brefeldin A (Sigma) for 4-5 hr before staining for flow cytometric analysis. Alternatively, cells were stimulated with 50 ng/mL PMA and 500 ng/mL ionomycin (Sigma) and 10 mM Brefeldin A for 4 hr before staining for flow cytometric analysis. Similarly, Small intestinal lamina propria cells were stimulated with 20 ng/mL IL-23, IL-6, IL-2 and IL-1β in the presence of 10 μM Brefeldin A for 2 hours, before PMA (50 ng/mL) and ionomycin (500 ng/mL) was added for a further 3 hours.

For co-culture of WT and CD200R1KO cells, dorsal skin cells were isolated and either the WT or CD200R1KO cells were labelled with 10 μM eF450-conjugated cell proliferation dye (Fischer Scientific) for 10 minutes in the dark at 37°C then were washed and co-cultured with the unlabelled cells.

#### ELISA

Levels of IL-17A, IL-10, IFN-γ, and IL-25 were measured using Ready-SET-Go! ELISA kits (eBioscience) following the manufacturer’s instructions.

#### Flow cytometric cell sorting

Cells were isolated from the small intestinal lamina propria and labelled with fluorescent antibodies as detailed above, before sorting on a BD FACS Aria cell sorter to a typical purity of >95%.

#### RNAseq

ILC3s (Live, single cell, CD45^+^, Lin^-^ (Ter119, F4/80, CD11b, CD11c, FceRIa, Gr1, CD19), CD3^-^, TCRb^-^, TCRgd^-^, CD90.2^+^, CD127^+^, GFP^+^) were sorted from RORc eGFP (WT) and CD200R1KO RORc eGFP (KO) small intestinal lamina propria keeping cells from each mouse separate. RNA was isolated using the RNeasy Micro kit (QIAGEN) following the manufacturer’s instructions. Contaminating genomic DNA was removed using gDNA Eliminator columns. Quality and integrity of the RNA samples were assessed using a 2200 TapeStation (Agilent Technologies, Cheadle, UK) and libraries were generated using the TruSeq^®^ Stranded mRNA assay (Illumina, Inc., California, USA) according to the manufacturer’s protocol before sequencing with an Illumina HiSeq4000 instrument. The output data was demultiplexed (allowing one mismatch) and BCL-to-Fastq conversion was performed using Illumina’s bcl2fastq software, version 2.17.1.14. Reads were quality trimmed using Trimmomatic_0.36 (PMID: 24695404) then mapped against the reference mouse genome, version mm10/GRCm38 using STAR_2.4.2 (PMID: 23104886). Counts per gene were calculated with HTSeq_0.6.1 (PMID:25260700) using annotation from GENCODE M12 (http://www.gencodegenes.org/). Normalisation and differential expression was calculated with DESeq2_1.12.3 (PMID:25516281) and edgeR_3.12.1 (PMID:19910308). Ingenuity Pathways Analysis was used to determine differentially regulated pathways.

#### Generating bone marrow chimeric mice

Male WT (C57BL/6) and CD200R1KO mice (both CD45.2) were exposed to a split dose of 10.5-11Gy irradiation. Bone marrow was isolated from male WT (CD45.1 or CD45.1 x C57BL/6 F1) and CD200R1KO (either CD45.1 or CD45.1 x CD200R1KO F1) mice and donor T cells were removed using CD90.2 microbeads and Miltenyi LS MACS columns. Cells were counted and mixed at a 1:1 ratio before being adoptively transferred intravenously into the irradiated hosts. 10-12 weeks post reconstitution, skin inflammation was induced by topical Aldara cream application daily (as described above) for 6 days. A day after the final treatment, mice were euthanised and cells analysed by flow cytometry.

#### Statistics

Data were analysed for normality by Shapiro-Wilk test. When comparing two independent groups a Student’s t test (for normally distributed data) was used. For three or more groups, data were analysed by One-way ANOVA (as normally distributed) or to determine the effects of two independent variables on multiple groups a Two-way ANOVA was used followed by a Bonferroni post hoc test. All statistical tests were performed using Prism Software (GraphPad Software Inc., USA). Values of less than p<0.05 were considered significant. Error bars show standard deviation.

All experiments were performed at least twice (except for the RNAseq analysis), with at least 3 independent samples per group. Each data point represents an individual animal.

## Abbreviations

CHILP: common helper-like innate lymphoid progenitor
DETC: dendritic epidermal T cell
IL: interleukin
ILC: innate lymphoid cell
ILC3s: group 3 innate lymphoid cells
Lti: lymphoid tissue inducer cell
NCR: Natural cytotoxicity receptor
ROS: Reactive oxygen species
TLR7: Toll-like receptor 7
Th17: Type 17 T helper cell

## Acknowledgments

This research was funded by a pre-competitive, open innovation award to the Manchester Collaborative Centre for Inflammation Research, by University of Manchester, AstraZeneca and GSK, and a Wellcome Trust and Royal Society, Sir Henry Dale Fellowship to AS (109375/Z/15/Z). The University of Manchester Bioimaging Facility microscopes used in this study were purchased with grants from BBSRC, Wellcome and the University of Manchester Strategic Fund. We acknowledge assistance from Peter Walker, Roger Meadows and Gareth Howell, and Leo Zeef and the use of the University of Manchester Histology, Flow Cytometry, Genomic Technologies, Bioinformatics and Biological Services facilities. We also acknowledge technical help from Peter Warn and Andrew Sharp on skin infection models and Tovah Shaw on intestinal cell isolation. We acknowledge intellectual expertise on ILC3s from Matthew Hepworth and on CD200R1 from Tracy Hussell.

## Data availability statement

The datasets generated during this study are available from the corresponding author on reasonable request.

## Author contributions

HL, SJ, LB, JC, MP and AS performed experiments. HL, AO, SJ and AS analyzed the data. AS designed the project and wrote the manuscript.

