## Supplementary information for "CD200R1 promotes IL-17 production by ILC3s, by enhancing STAT3 activation"

**Supplementary methods**

**Skin infection model**

*C. albicans* SC5314 (provided by Dr Andrew Sharp) was grown on Sabouraud agar (SAB) plates supplemented with 50 μg/mL chloramphenicol (Fischer Scientific) overnight. Colonies were cultured for 18-24 hours at 37°C in yeast extract-peptone-dextrose (YPD) broth (Fischer Scientific). The fungi were washed with PBS and 2 x $10^7$ CFU was applied topically to shaved dorsal skin. After 48hr mice were euthanised and analysed. Fungal burden was determined by lysing small skin pieces in Precellys hard tissue lysing tubes (Stretton Scientific) containing PBS and 1% penicillin and streptomycin, using a Precellys 24 homogenising instrument (Stretton Scientific). Lysate was cultured on YPD plates supplemented with 50 μg/mL chloramphenicol at 30°C for 24 hours before colony enumeration.

**Supplementary Table 1**

Antibodies used for flow cytometry

| **Antibody specificity** | **Clone** | **Manufacturer** |
|---|---|---|
| B220 | RA3/6B2 | eBioscience |
| CCR2 | 475301 | R & D systems |
| CCR6 | 29-2L17 | Biolegend |
| CD1d | 1B1 | Biolegend |
| CD3ε | 145-2C11 | eBioscience |
| CD4 | RM4-5 | Biolegend |
| CD11b | M1/70 | eBioscience |
| CD11c | N418 | eBioscience |
| CD19 | eBio1D3 | eBioscience |
| CD24 | M1/69 | Biolegend |
| CD25 | PC61 | Biolegend |
| CD45 | 30-F11 | eBioscience |
| CD45.1 | NDS58 | BD Bioscience |
| CD45.2 | 104 | eBioscience |
| CD64 | X54-5/7.1 | Biolegend |
| CD90.2 | 53-2.1 | eBioscience |
| CD117 (cKit) | 2B8 | eBioscience |
| CD127 | A7R34 | eBioscience |
| CD200R1 | OX110 | eBioscience |
| CD207 | eBioL31 | eBioscience |
| F4/80 | BM8 | eBioscience |
| FcεRIα | MAR-1 | eBioscience |
| GATA3 | TWAJ | eBioscience |
| Gr1 (Ly-6G/Ly-6C) | RB6-8C5 | eBioscience |
| IL-17A | eBio17B7 | eBioscience |
| IL-23R | 3C9 | BD Biosciences |
| Ly6C | HK1.4 | eBioscience |
| Ly-6G | 1A8 | BD Biosciences |
| MHCII (IA-IE) | M5/114.15.2 | eBioscience |
| NKp46 | 29A1.4 | eBioscience |
| RORγt | B2d | eBioscience |
| ScaI (Ly6A/E) | D7 | Biolegend |
| TCRβ | H57-597 | BD Bioscience |
| TCRγδ | eBioGL3 | eBioscience |
| Ter119 | TER-119 | eBioscience |

**Figure S1: CD200R1 deficiency reduces the IL-17A production by ILCs in response to *C. albicans* skin infection**

WT and CD200R1KO (KO) mouse dorsal skin was infected with $10^7$ CFU *C. albicans SC5314* (or PBS) and after 2 days mice were euthanised and disease severity was analysed. **A.** Skin thickening measured using a digital micrometer. **B.** H&E stained skin sections. **C.** Fungal burden in skin. **D.** Proportion of live single cells that are $CD45^+$ in infected skin. **E.** Proportion of $CD45^+$ cells that are neutrophils in infected skin. **F.** Proportion of skin $CD3^{low}$ $\gamma\delta$ T cells producing IL-17A. **G.** Proportion of skin ILCs producing IL-17A. Data shown are from 2 pooled independent experiments. A, D-G Data were analysed by two-way ANOVA followed by a Bonferroni post hoc test. C data were analysed by Student's t test.

**Figure S2: Known ILC3 inhibitory cytokines are not increased in CD200R1KO skin cell cultures**

Total dorsal skin cells were isolated from WT and CD200R1KO (KO) mouse dorsal skin and were stimulated with IL-23 overnight. IL-10, IFN$\gamma$ and IL-25 levels were measured in culture supernatant by ELISA. Data are from one experiment, representative of two independent experiments. Data were analysed by Student's t test.

**Figure S3: Flow cytometric sorting of Rorc eGFP+ ILC3s**

Small intestinal lamina propria cells were isolated, stained with antibodies and were sorted using a flow cytometer. **A.** Cells prior to flow cytometric cell sorting and **B.** Purity of the sorted cells ($CD45^+$ $Lineage^-$ $Thy1^+$ $CD127^+$ RORc $eGFP^+$). Data shows representative flow cytometric sorting gating and purity.

**Figure S1**

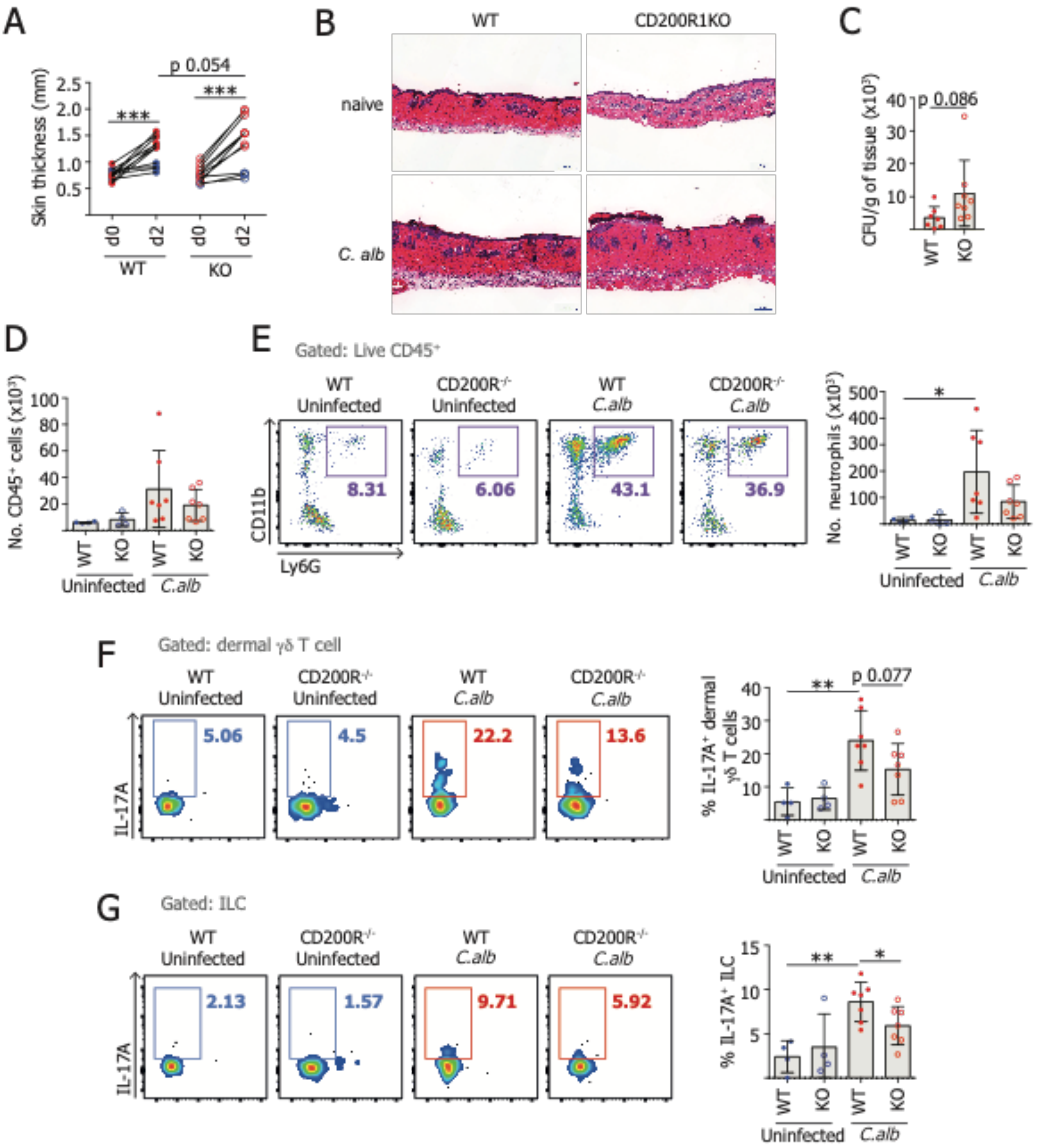

A
p 0.054
***
***
Skin thickness (mm)
d0
d2
d0
d2
WT
KO
B
WT
CD200R1KO
naive
C. alb
C
CFU/g of tissue (x10³)
p 0.086
WT
KO
D
No. CD45⁺ cells (x10³)
WT
KO
WT
KO
Uninfected
C.alb
E
Gated: Live CD45⁺
WT Uninfected
CD200R⁻/⁻ Uninfected
WT C.alb
CD200R⁻/⁻ C.alb
CD11b
Ly6G
8.31
6.06
43.1
36.9
No. neutrophils (x10³)
*
Uninfected
C.alb
F
Gated: dermal γδ T cell
IL-17A
5.06
4.5
22.2
13.6
% IL-17A⁺ dermal γδ T cells
**
p 0.077
G
Gated: ILC
2.13
1.57
9.71
5.92
% IL-17A⁺ ILC
**
*

**Figure S2**

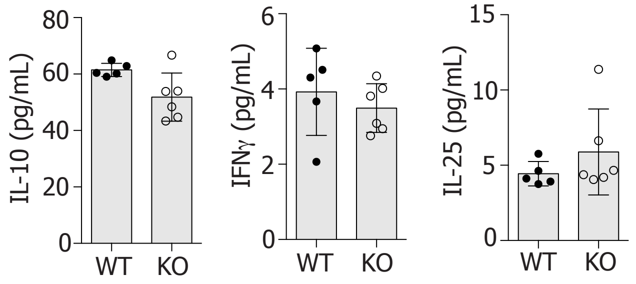


**Figure S3**

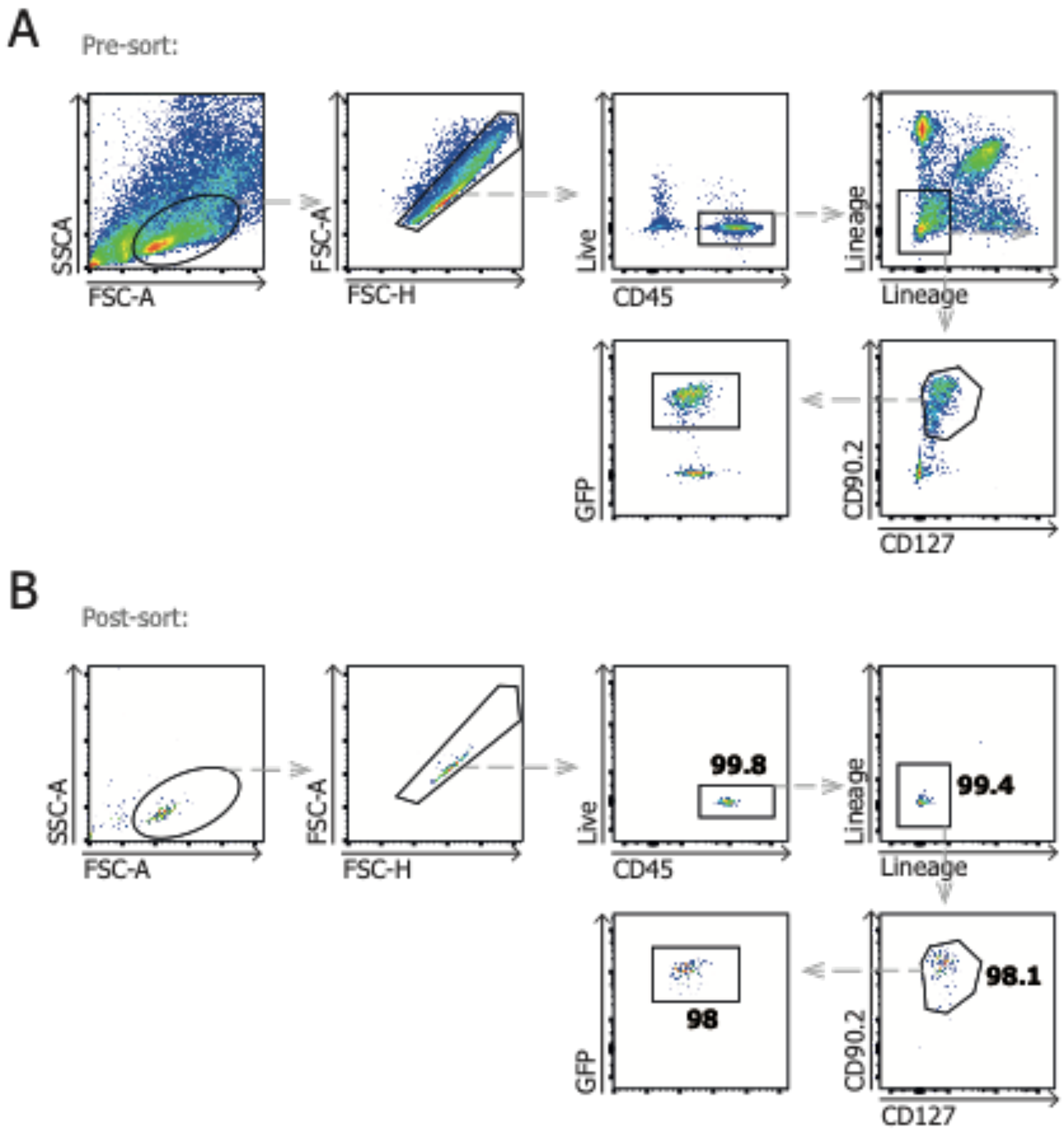
